# Salivary Glands treated with AAV vectors encoding PYY and exendin-4 impact upon taste perception and body mass accumulation in mice

**DOI:** 10.1101/2022.12.06.519331

**Authors:** S. Iyer, JP Montmayeur, S. Zolotukhin, C.D. Dotson

## Abstract

The availability of high-calorie foods is likely a causative factor for high rates of obesity and metabolic disorders, which have been linked to food intake dysregulation. Several gut peptides have been implicated in feeding modulation and body mass accumulation. For example, glucagon peptide-like 1 (GLP-1) and peptide tyrosine-tyrosine (PYY) have been shown to mediate satiety and reduce food intake. While systemic administration of such peptides has been explored as a therapy for metabolic disease, the effects of these hormones on taste signaling should also be considered given the importance of taste to feeding decisions. Peptide signaling systems are present in taste buds and oral GLP-1 and PYY signaling has been shown to influence taste responsiveness and feeding. Indeed, we previously demonstrated that genetic knockout of PYY in mice can impact on taste responsiveness and feeding and that viral overexpression of PYY in the salivary glands of these mice can rescue responsiveness. The present work uses AAV-mediated salivary gland treatment of both GLP-1 receptor agonist exendin-4 and PYY encoding vectors to explore the impact of the presence of these peptides on taste and body-mass accumulation in wild-type mice with intact peptide signaling systems. Results showed a significant effect of salivary gland treatment on responsiveness to multiple taste qualities. Treatment with a vector designed to overexpress both peptides in saliva resulted in substantial reduction in body mass accumulation. These findings show taste modulation and impacts on body mass accumulation by the targeting of salivary glands with vectors designed to overexpress metabolic peptides in wild-type mice and suggest that the taste bud is a promising substrate for food intake modulation.

## INTRODUCTION

The desire for good tasting foods spans all ages, races, and cultures (Behrens, Meyerhof, Hellfritsch, & Hofmann, 2011; Drewnowski, 1997a, 1997b; Drewnowski, Mennella, Johnson, & Bellisle, 2012). Highly appetitive foods are often rich in carbohydrates and/or fats and a preference for these energy sources can orient feeding behavior and, therefore, promote survival (Drewnowski & Almiron-Roig, 2010; Drewnowski, et al., 2012). However, in our current environment, the ubiquitous availability of high-calorie foods is likely one of the causative factors behind the alarming rise in obesity rates, obesity-related conditions (Austin, Ogden, & Hill, 2011; Centers for Disease & Prevention, 2004; Ford & Dietz, 2013), and the resulting decrease in lifespan for obese individuals (Biener, Cawley, & Meyerhoefer, 2017). The cost of this public health crisis has been staggering: in the year 2013, it was estimated that healthcare costs associated with obesity totaled ~$342 billion, with medical costs for obese individuals ~$3,500 higher than for those of normal weight (Biener, et al., 2017; Finkelstein, Trogdon, Cohen, & Dietz, 2009).

Obesity and related metabolic disorders have been linked to the dysregulation of food intake (Garaulet & Gómez-Abellán, 2014; Narayanaswami & Dwoskin, 2017). Several peptide hormones secreted by the gut in response to the presence or absence of food have been shown to modulate feeding behavior. For example, the orexigenic peptide ghrelin secreted in the absence of food by primary gastric cells has been shown to enhance food intake (Gil-Campos, Aguilera, Canete, & Gil, 2006). Conversely, glucagon-like peptide 1 (GLP-1) and peptide YY (PYY) are secreted by enteroendocrine L cells in response to food and have been shown to mediate satiety and reduce food intake (Kadouh, et al., 2020b; Kjaergaard, et al., 2019; Reidelberger, Haver, & Chelikani, 2013; Shi, et al., 2012; Yang, Moghadam, Cordner, Liang, & Moran, 2014b). The influence of these peptides on food intake and thus body mass accumulation underscores the potential value of such hormones as treatments for obesity and related metabolic disease. Indeed, this potential has led to the utilization of these peptides as treatments for various diseases. Glucagon therapy is well established in the treatment of hypoglycaemia (Pearson, 2008), while GLP-2 analogues are currently used in the treatment of short bowel syndrome (Hargrove, et al., 2020; Jeppesen, et al., 2011; Jeppesen, et al., 2005). Furthermore, since approval of the first GLP-1 mimetic for the management of Type 2 diabetes mellitus (T2DM), GLP-1 therapeutics have become the gold standard in the treatment of T2DM (Edwards, et al., 2001; Lafferty, O’Harte, Irwin, Gault, & Flatt, 2021; Wilkinson, et al., 2018).

The concept of multi-purpose peptide drugs, efficacious for the simultaneous treatment of several disease states, has become the focus of intense biomedical research in recent years. These designer peptides have expanded biological functions owing to their multi-receptor specificity (Chepurny, et al., 2018; Kim, Lin, Blomain, & Waldman, 2014). Indeed, new multi-agonist peptides have been designed so that they have the capacity to stimulate different receptor systems simultaneously (Dalboge, et al., 2014; Frias, et al., 2017; Sanchez-Garrido, et al., 2017) and are currently being evaluated as possible treatments for a variety of metabolic conditions including obesity and T2DM (De Silva, et al., 2011; Finan, et al., 2015; Frias, et al., 2017; Sanchez-Garrido, et al., 2017; Trevaskis, et al., 2013).

One of the many factors influencing food intake that has not yet been targeted by the pharmaceutical industry is sensory information/processing. A large amount of sensory information from both the oronasal cavity and gastrointestinal tract is sent to the central nervous system (CNS), imparting information related to macronutrient composition, caloric density, osmolarity, and potential toxicity of food (Ahima & Antwi, 2008; Concas, et al., 2022; Morrison & Berthoud, 2007; Saper, Chou, & Elmquist, 2002; Zheng, Lenard, Shin, & Berthoud, 2009). Indeed, it is the sense of taste which acts to protect the rest of the alimentary canal by providing information on which nutrients to ingest and which to reject (Concas, et al., 2022; Forde & de Graaf, 2022; Mattes, 2003; Scott & Verhagen, 2000; Tanaka, Reed, & Ordovas, 2007).

Interestingly, in addition to their presence in the GI tract and CNS, both GLP-1 and PYY, as well as their cognate receptors, are expressed in cells of the peripheral gustatory system (Acosta, et al., 2011b; Hurtado, et al., 2012; Hurtado, et al., 2013; B. Martin, et al., 2009; Shin, et al., 2008). We have shown that PYY is also present in the saliva of both humans and mice and that the augmentation of salivary PYY_3-36_, using either genetic and/or pharmacological approaches, affects taste responsiveness (La Sala, et al., 2013), as well as reduces food intake and body weight in diet-induced obese mice (Acosta, et al., 2011a). Oral GLP-1 signaling also impacts upon taste responsiveness (e.g., Brindisi, et al., 2019; Kadouh, et al., 2020a; B. Martin, et al., 2009; C. Martin, et al., 2012; Shin, et al., 2008). Together, these findings indicate that taste function is subject to modulation by metabolic hormones, and that it is possible that these functions can be experimentally exploited to achieve long-term therapeutic effects.

The use of multi-receptor system peptides have shown promise in the treatment of various disorders (see Lafferty, et al., 2021 for a review). However, it is unclear whether the simultaneously stimulation of multiple receptor systems can impact on the functioning of the peripheral gustatory system. The present study explores whether AAV-mediated expression of both GLP-1 and PYY in the oral cavity, in wild-type mice with intact hormone signaling systems, can affect taste responsiveness and body mass accumulation.

## MATERIALS AND METHODS

This study was approved by the Institutional Animal Care and Use Committee (IACUC) at the University of Florida. All procedures in the study were carried out in accordance with the principles of the National Research Council’s guide for the care and use of laboratory animals. The mice were housed at 22-24°C in a twelve-hour darklight cycle with *ad libitum* access to water and food unless specified otherwise.

### Behavioral assays

#### Mice

All mice used in behavioral experiments were 10-12 weeks old. Vector treated C57BL/6J served as subjects for brief-access taste testing, as well as for the analysis of the impact of vector treatment on body mass accumulation. Mice were administered either AAV5-PYY-miR, AAV5-Ex-4-miR, AAV5-PYY+Ex-4-miR, or AAV5-GFP-miR 1 month before the first day of training. Exendin-4 is a 39 amino acid agonist of the GLP-1 receptor. It was selected because it has a significantly longer half-life than GLP-1 and is used in the treatment of T2DM (Ding, Saxena, Lin, Gupta, & Anania, 2006; Edwards, et al., 2001; Egan, Clocquet, & Elahi, 2002; Wilkinson, et al., 2018). AAV was administered by submandibular salivary gland cannulation as described previously (Crosson, et al., 2019; Katano, et al., 2006; La Sala, et al., 2013). After vector administration, mice were single housed and given *ad libitum* access to food and water until the start of the training/testing sessions.

In conditioned taste aversion (CTA) experiments, PYY KO mice and wild-type (WT) controls generated at the Garvan Institute of Medical Research as described previously (Boey, et al., 2006) served as subjects. All mice were housed individually in standard cages with bedding. Mice were habituated to their environment for at least seven days before testing began.

#### Taste Stimuli

Tastants were prepared with purified water (Elix 10; Millipore, Billerica, MA, USA) and reagent grade chemicals. Tastants were presented to the mice at room temperature. Presented tastants include sucrose, intralipid, NaCl, quinine hydrochloride, and citric acid. Testing stimuli consisted of a “no stimulus” water control and 5 concentrations of each tastant: sucrose (31.25, 62.5, 125, 250, and 500 mM; Fisher Scientific, Atlanta, GA, USA); intralipid (1.25, 2.5, 5, 10, and 20%; Baxter Healthcare, Deerfield, IL, USA); NaCl (30, 100, 300, 600, and 1000 mM; MilliporeSigma, Burlington, MA, USA); quinine hydrochloride (DB; 0.03, 0.1, 0.3, and 1, 3 mM; MilliporeSigma); and citric acid (CA; 1, 3, 10, 30, and 100 mM; MilliporeSigma).

For CTA experiments, the condition stimulus (CS) was 1% linoleic acid (LA; MilliporeSigma). The test stimuli (TS) were 0.5, 1, and 2% linoleic acid, as well as 5% intralipid.

#### Brief-Access Taste Testing

Brief-access taste testing was administered within a Davis Rig Gustometer (Davis MS-160; DiLog Instruments, Tallahassee, FL, USA; Smith, 2001) as previously described (e.g., Crosson, et al., 2019; Dotson, Roper, & Spector, 2005; Dotson & Spector, 2004, 2005, 2007; Elson, Dotson, Egan, & Munger, 2010; La Sala, et al., 2013; Shin, et al., 2008). Two testing protocols were used – one for preferred tastants and one for aversive tastants. Testing consisted of 25-min sessions during which mice were presented access to the sipper tubes for 5 sec with 7.5 sec inter-presentation intervals.

For appetitive stimuli (i.e., sucrose and intralipid), the animals were first trained to lick a stationary tube of water for 30 min in the Davis rig after being placed on ~23.5 h restricted water access schedule. Animals then received 2 days of testing with one of the appetitive stimulus arrays and purified water while maintained on the water-restriction schedule. This was done to familiarize the animals with the testing procedure and the stimulus array. The mice were subsequently tested for three consecutive days. During testing, mice were restricted to 1 g of food and 2 ml of water for 23.5 hours prior to testing.

For aversive stimuli, animals were trained to lick a stationary tube of water for 30 min in the Davis rig after being placed on ~23.5 h restricted water access schedule. Animals then received 2 days of testing with a ‘water-only’ stimulus array while maintained on the water-restriction schedule. This was done to familiarize the animals with the testing procedure. The mice were subsequently tested for three consecutive days with one of the aversive stimulus arrays detailed above and purified water. Following presentation of an aversive tastant, mice were presented with a 1 sec H_2_O rinse to minimize crossing over effects.

If mice dropped below 85% of their starting body weight, they received 1 ml of water after the testing session was completed.

#### Statistical Analysis – vector-treated mice

For the normally avoided stimuli, the average number of licks per trial for each concentration was divided by that animal’s average licks per trial to water yielding a tastant/water lick ratio (Glendinning, Gresack, & Spector, 2002). To control for the low rate of water licking when animals are tested with normally preferred stimuli, a “tastant minus licks to water” difference score was also derived by taking the mean number of licks to water and subtracting it from the mean number of licks at each concentration (Jiang, Blonde, Garcea, & Spector, 2008; Malone, et al., 2021; Spector, Redman, & Garcea, 1996; Treesukosol, Blonde, & Spector, 2009). All scores were analyzed with analyses of variance (ANOVAs). The main variable of interest was vector treatment type (i.e., each experimental vector group was compared directly to B6 controls treated with a GFP expressing vector). If a main effect was not observed, a vector treatment type X concentration interaction was explored. If a significant interaction was observed, *post hoc* t tests were conducted to determine which concentrations differed between the experimental groups. The conventional statistic p ≤ 0.05 was applied as the statistical rejection criterion. Only mice that had at least one trial at every concentration were included in the analysis of a given stimulus. One mouse treated with a vector encoding PYY was excluded from the analysis of quinine hydrochloride responsiveness. Lastly, If a main effect of vector treatment type was observed, t tests were conducted to determine if the groups also differed in the average number of test trials taken over the experimental period.

For the analysis of body mass accumulation, the main variable of interest was again vector treatment type. If a main effect was not observed, a vector treatment type X week interaction was explored. If a significant interaction was observed, *post hoc* t tests were conducted to ascertain which weeks did body weight differed between the experimental groups. The conventional statistic p ≤ 0.05 was again applied as the statistical rejection criterion.

### Plasmid construction

To restrict the leaky expression of the transgene in the liver or skeletal muscle, we utilized triplicates of miR122 and miR206 target sites incorporated into the 3’-UTR of the transgene cassettes (**Fig. 1**), as described earlier (Crosson et al, 2019). We have constructed the following vectors: pTR-GFP-PYY-miRNA, pTR-GFP-Ex-4-miR, pTR-PYY-Ex-4-miR, and pTR-GFP-miR (negative control vector).All plasmid constructs were validated by sequencing.

**Figure 1.**
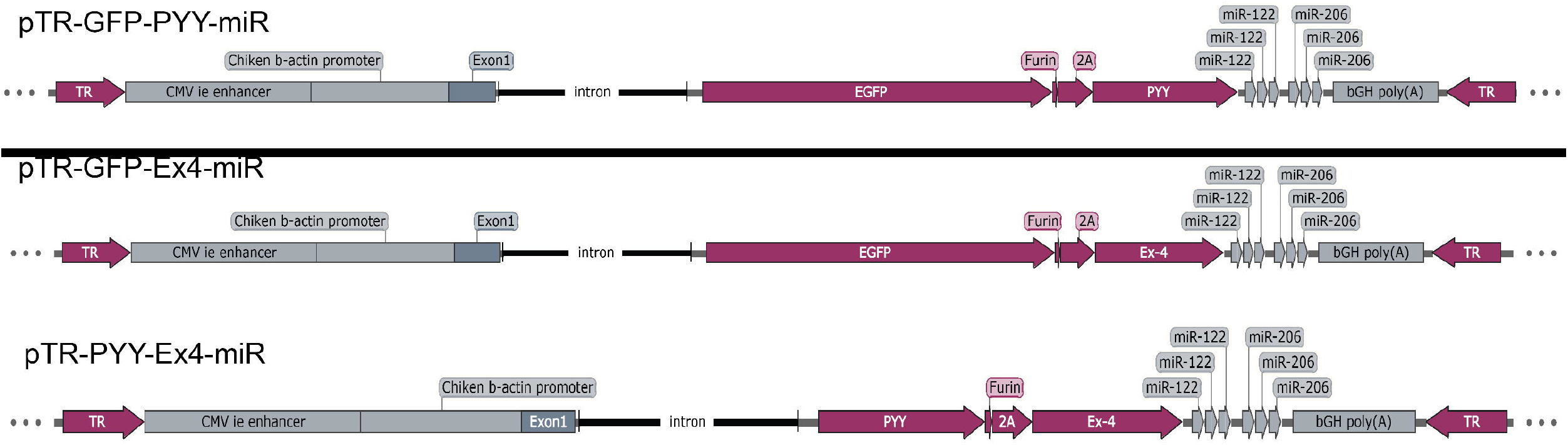
Genetic maps of rAAV vectors used in the study. Please see Methods section for details.

### AAV vector production

Recombinant AAV vectors were produced in HEK 293 cells using a triple plasmid-based transfection method and purified via Iodixanol density centrifugation as described previously (Zolotukhin, et al., 1999). pHelper (Agilent cat no. 240071-52) was used to supply the adenoviral helper genes and pACG2R5C (Zolotukhin, et al., 2002) was used to supply the AAV2 rep and AAV5 cap genes. Vectors were titered using a PicoGreen based assay described by Piedra et al. (2015), and were sterile filtered before administration to animals.

#### Conditioned Taste Aversion Testing

All mice were 10 – 12 weeks of age at the time of testing. These procedures have been detailed previously (Dotson & Spector, 2007). In brief, mice were first trained to behave in Davis rig gustometers. Water bottles were removed 24 hours prior to testing. Mice were trained to lick in the Davis rig for 2 days, ad libitum to vehicle (i.e., 0.3% xanthum gum) during 30 min sessions. Mice were then trained to lick vehicle, in 5-second trials, for an additional 3 days.

After this training, mice were then conditioned. Water bottles were removed 24 hours prior to conditioning. On days 1, 2, 4, 5, 7 and 8 of conditioning, vehicle intake was measured during a morning 15-min test in the animal’s home cage. On days 3, 6, and 9 of conditioning, the CS (i.e., 1% LA) was given during the morning of the 15-min test immediately followed by an injection of 0.15 M LiCl or saline (3.0 mEq/kg BW). Food was removed during the 15-min test on all days. Approximately 5 hours after each conditioning session, a 1-hour dH2O intake session was given in the afternoon (food accessible at this time).

After the conditioning phase of the experiment, the mice were tested for their responsiveness to the test stimulus in the Davis rig. Mice were water-deprived approximately 23.5 hours prior to testing. Mice were tested for 1 day with vehicle in 5-second trials. Then mice were tested for 1 day with the TS array.

#### Statistical Analysis – CTA

Tastant/vehicle lick ratios were calculated for each mouse representing the mean number of test stimulus licks per trial divided by the mean number of vehicle licks per trial (vehicle rinse trials were not included in the analysis). All of these data were statistically examined with ANOVAs and t tests. The statistical rejection criterion was set at the conventional value of p ≤ 0.05.

## RESULTS

Previously, we found that PYY germline KOs displayed impaired behavioral responsiveness towards a bitter tasing stimulus, as well as towards the fatty acid emulsion, intralipid (La Sala, et al., 2013). Moreover, when PYY levels were augmented in saliva of PYY KO mice, behavioral responsiveness to intralipid is rescued. We also showed that GLP-1R germline KOs showed decreased responsiveness to sucrose (B. Martin, et al., 2009; Shin, et al., 2008). These data suggest that manipulating oral peptide signaling can be exploited to impact on taste-related behavior and, potentially on body mass accumulation. In the current report, we asked whether treatment with vectors encoding these or related peptides in the salivary glands of WT mice, which have intact oral peptide signaling systems, would impact on taste-related behavior and body mass accumulation.

The treatment of salivary glands with vectors encoding either PYY, exendin-4, or both did not impact on the responsiveness of WT mice towards sucrose in either the deprived [all *Fs*(1,14) ≤ 0.98, *Ps* ≥ 0.34; **Fig. 2A**] or food and water-restricted conditions [all *Fs*(1,14) ≤ 0.38, *Ps* ≥ 0.55; **Fig. 2B**]. However, when compared to mice treated with a vector encoding GFP, those treated with vectors encoding both PYY and exendin-4 were hyperresponsive to intralipid in both the deprived [*F*(1,14) **=** 6.18, *P* = 0.03; **Fig. 3A**] and food and water-restricted conditions [*F*(1,14) **=** 10.99, *P* = 0.005; **Fig. 3B**]. These vector treated mice also initiated more trials over the testing period, relative to control mice, but only in the food and water-restricted condition [*t*(14) = 2.46, *P* = 0.03].

**Figure 2.**
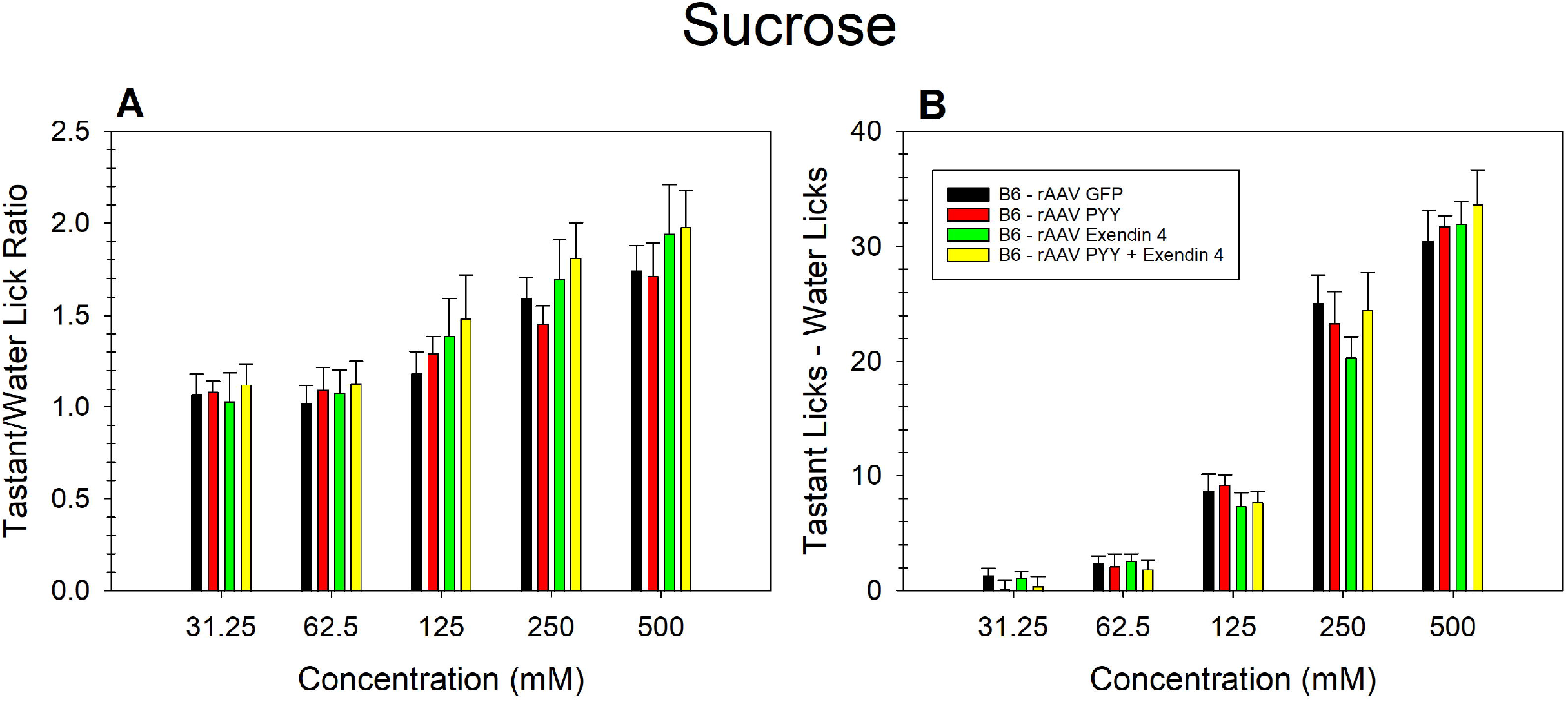
The treatment of salivary glands with vectors designed to overexpress either PYY, exendin-4, or both did not impact on the responsiveness of WT B6 mice towards sucrose in either the deprived or food and water-restricted conditions relative to vector treated GFP mice. Brief-access taste testing of WT mice treated with a vector encoding GFP (n = 8; black bars); WT mice treated with a vector encoding PYY (n =8; gray bars with right hashes); WT mice treated with a vector encoding exendin-4 (n = 8; dark gray bars with left hashes); and WT mice treated with a vector encoding both exendin-4 and PYY (n = 8; light gray bars with double hashes) in response to **(A)** sucrose (after water deprivation; all *Ps* ≥ 0.34), **(B)** sucrose (after food and water restriction; all *Ps* ≥ 0.55). For mice sampling sucrose after food and water restriction, a “tastant minus licks to water” difference score was derived by taking the mean number of licks to water and subtracting it from the mean number of licks at each concentration. For all other conditions and stimuli, the average number of licks per trial for each concentration was divided by that animal’s average licks per trial to water yielding a tastant/water lick ratio. Data are presented ± SEM.

**Figure 3.**
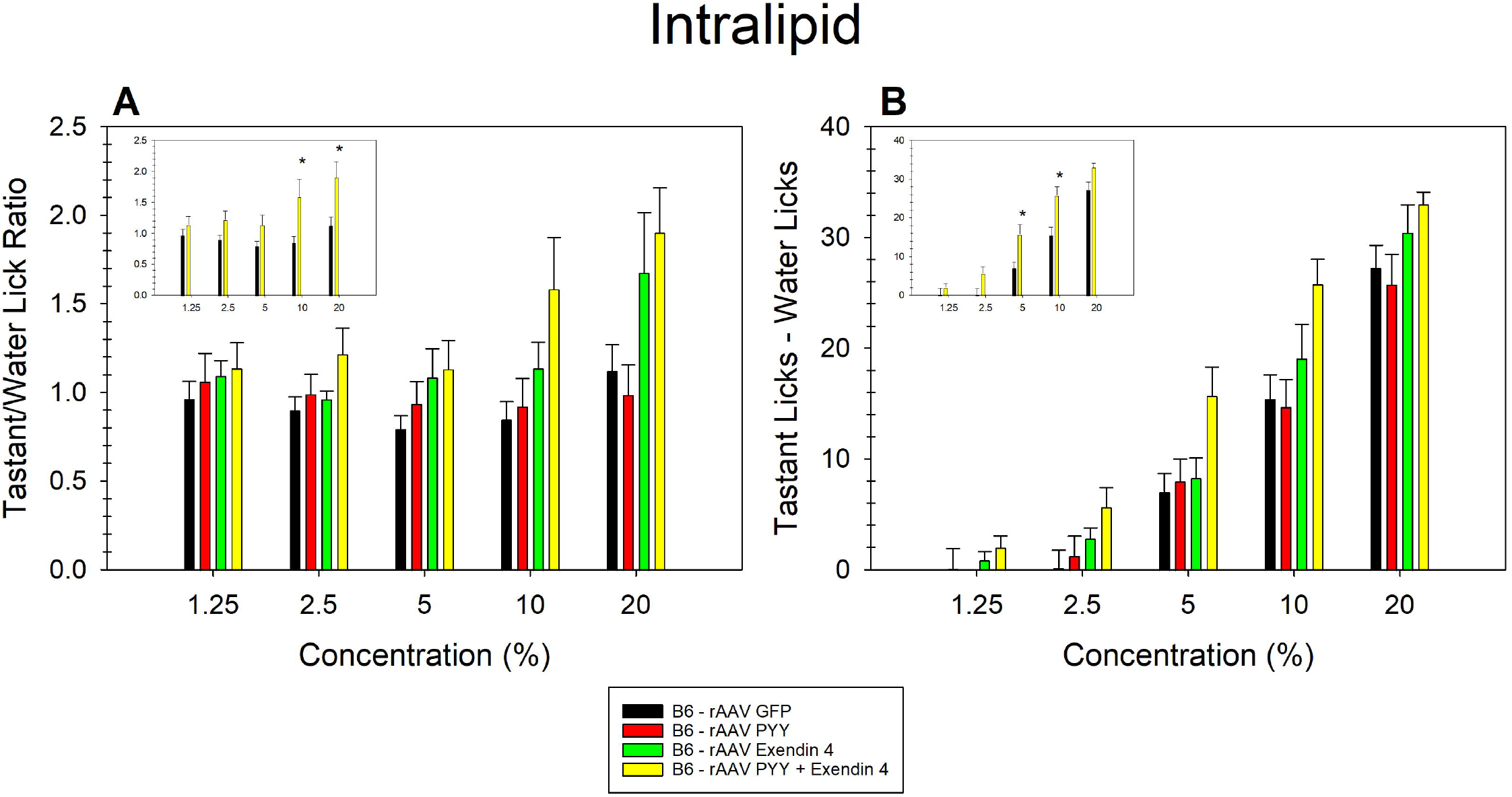
WT mice treated with a vector encoding both PYY and exendin-4 were hyperresponsive to intralipid, in both the deprived and food and water-restricted conditions, relative to WT mice treated with a vector encoding GFP. Brief-access taste testing of WT mice treated with a vector encoding GFP (n = 8; black bars); WT mice treated with a vector encoding PYY (n = 8; gray bars with right hashes); WT mice treated with a vector encoding exendin-4 (n = 8; dark gray bars with left hashes); and WT mice treated with a vector encoding both exendin-4 and PYY (n = 8; light gray bars with double hashes) in response to **(A)** intralipid (after water deprivation), **(B)** intralipid (after food and water restriction). When compared to mice treated with a vector encoding GFP, mice treated with a vector encoding both PYY and exendin-4 were hyperresponsive to intralipid in both the deprived [*P* = 0.03] and food and water-restricted conditions [*P* = 0.005; see insets in panels **A** and **B**]. For mice sampling intralipid after food and water restriction, a “tastant minus licks to water” difference score was derived by taking the mean number of licks to water and subtracting it from the mean number of licks at each concentration. For all other conditions and stimuli, the average number of licks per trial for each concentration was divided by that animal’s average licks per trial to water yielding a tastant/water lick ratio. Data are presented ± SEM.

When tested with normally avoided stimuli, mice treated with a vector encoding PYY were hyperresponsive when sampling QHCL relative to GFP-treated mice [*F*(1,13) **=** 5.95, *P* = 0.03; **Fig. 4A**]. However, these mice did not initiate more trials over the testing period, relative to control mice [*P* = 0.59]. WT mice treated with a vector encoding exendin-4 were hyperresponsive when sampling NaCl relative to GFP-treated mice [*F*(1,14) **=** 12.06, *P* = 0.004; **Fig. 4B**). These mice also did not initiate more trials over the testing period, relative to control mice [*P* = 0.22]. Treatment with either PYY, exendin-4, or vectors that encode both peptides did not impact on the responsiveness of WT mice towards citric acid [all *Ps* ≥ 0.14; **Fig. 4C**].

**Figure 4.**
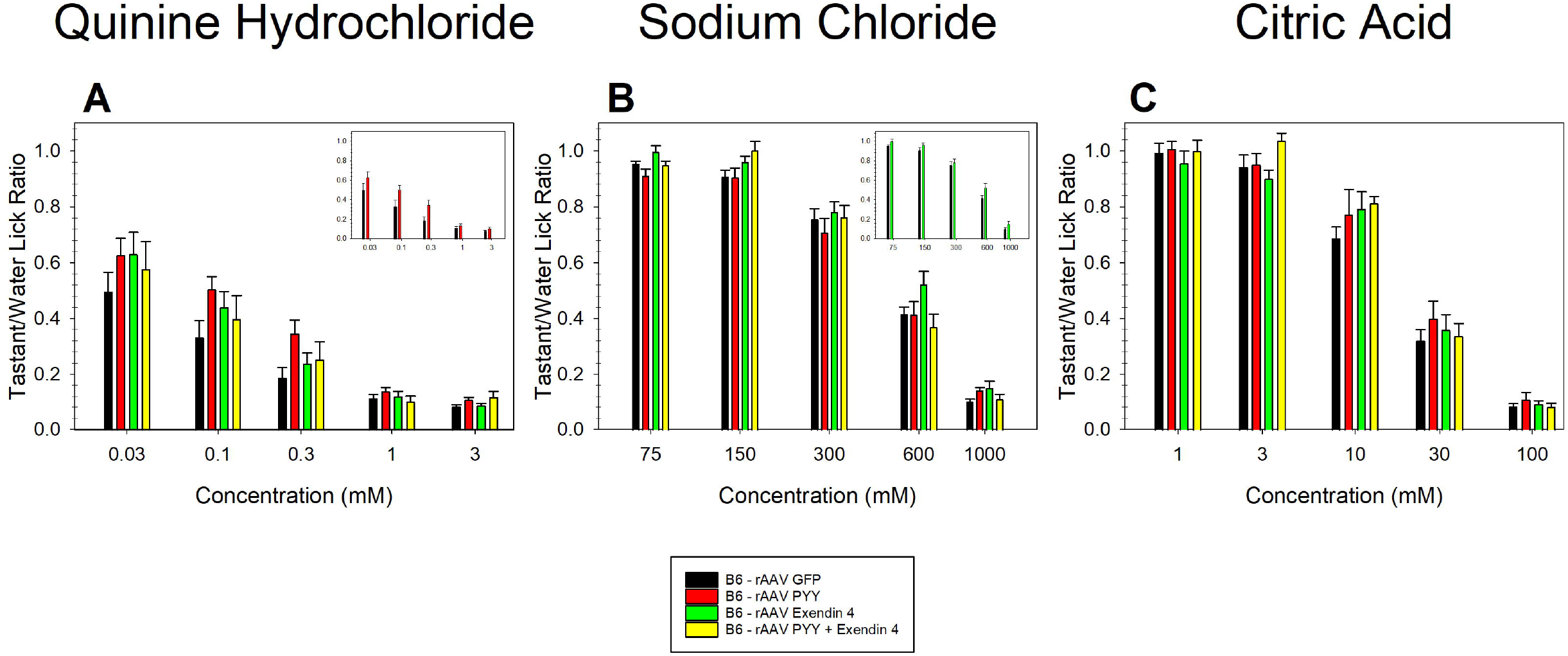
WT mice treated with a vector encoding PYY were hyperresponsive to quinine relative to WT mice treated with a vector encoding GFP and those treated with a vector encoding exendin-4 were slightly more responsive to sodium chloride relative controls. Brief-access taste testing of WT mice treated with a vector encoding GFP (n = 8; black bars); WT mice treated with a vector encoding PYY (n = 8; gray bars with right hashes); WT mice treated with a vector encoding exendin-4 (n = 8; dark gray bars with left hashes); and WT mice treated with a vector encoding both exendin-4 and PYY (n = 8; light gray bars with double hashes) in response to **(A)**quinine hydrochloride, **(B)**sodium chloride, and **(C)**citric acid. Mice treated with a vector encoding PYY were hyperresponsive when sampling QHCL relative to GFP-treated mice [*P* = 0.03; see inset in panel **A**] and mice treated with a vector encoding exendin-4 were hyperresponsive when sampling NaCl relative to control mice [*P* = 0.004; see inset in panel **B**]. Treatment with vectors encoding either PYY, exendin-4, or both did not impact on the responsiveness of WT mice towards citric acid [all *Ps* ≥ 0.14]. For all conditions and stimuli, the average number of licks per trial for each concentration was divided by that animal’s average licks per trial to water yielding a tastant/water lick ratio. Data are presented ± SEM.

To assess the impact of oral peptide expression on body mass accumulation, we measured the body mass of these mice once a week for 12 weeks **(Fig. 5)**. Exendin-4 treated mice gained slightly less weight relative to GFP-treated mice control over the 12-week test period [*F*(1,18) **=** 4.70, *P* = 0.04]. PYY-treated mice also gained slightly less weight than GFP-treated mice. However, this difference was only apparent during the latter weeks of testing [*F*(11,198) **=** 265.0, *P* = 0.0004, *interaction*]. Mice treated with a vector encoding both exendin-4 and PYY gained sustainably less weight relative to GFP-treated mice over the 12-week testing period [*F*(1,18) **=** 29.50, *P* = 0.00004]. *Post hoc* t-tests revealed that exendin-4 treated mice weighed significantly less than GFP-treated controls starting 6 weeks after testing and remained lighter across the remainder of the 12-week testing period. For PYY-treated mice, *post hoc* t-tests revealed that although the mice weighed less than GFP vector-treated controls on many of the latter weeks of testing, none of these differences reached statistical significance [all *Ps* ≥ 0.14]. Mice treated with a vector encoding both exendin-4 and PYY gained sustainably less weight starting 1 week after testing and remained lighter across the remainder of the 12-week testing period. Indeed, this difference was even more pronounced when the animals were switched to a high-fat diet at week 9.

**Figure 5.**
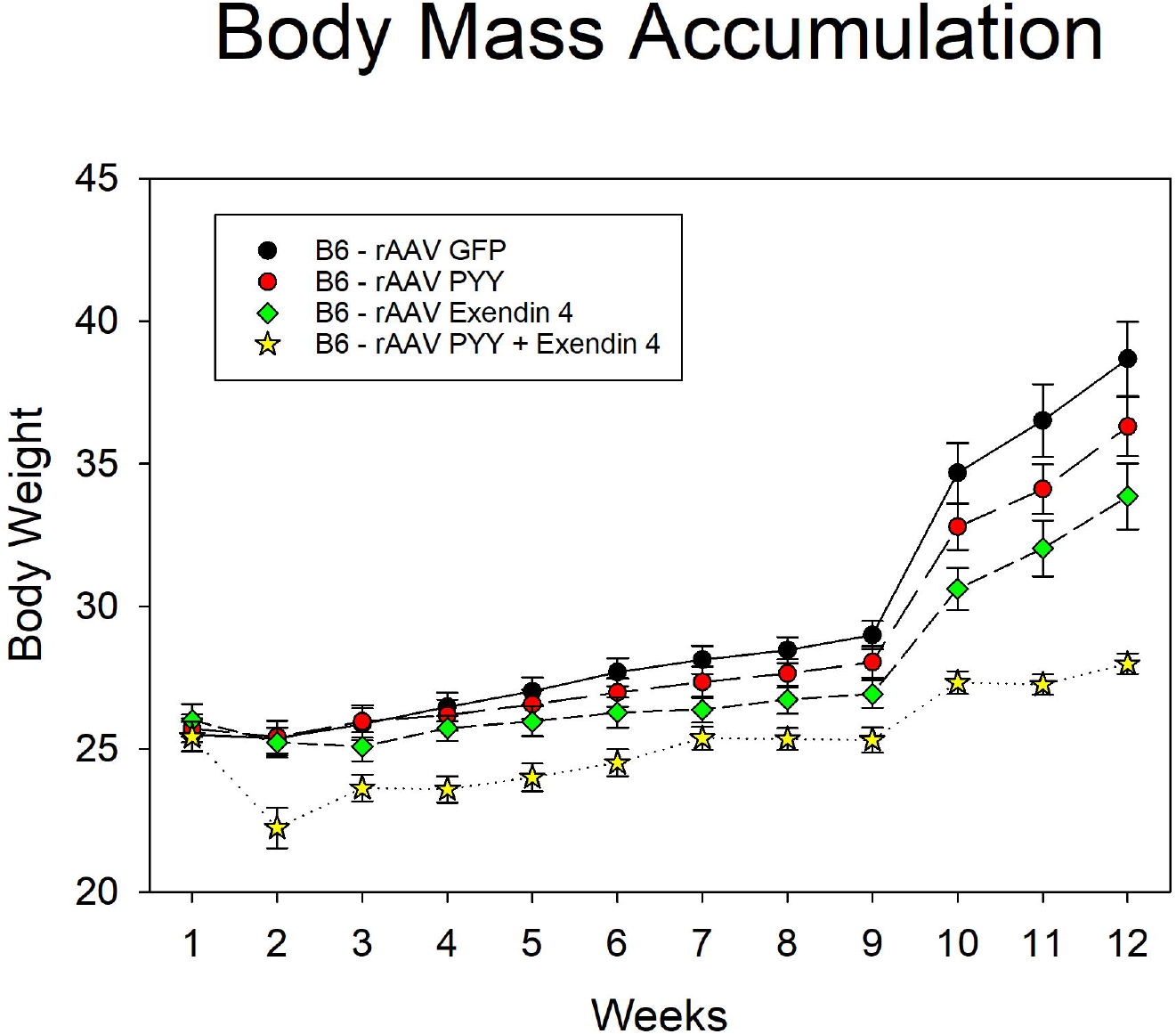
WT mice whose salivary glands were treated with vectors encoding metabolic peptides gained less body mass relative to mice treated with a vector encoding GFP. Body mass measurements of WT mice treated with a vector encoding exendin-4 (n = 10; open circles); WT mice treated with a vector encoding PYY (n = 10; gray diamonds); WT mice treated with a vector encoding both exendin-4 and PYY (n = 10; gray stars); and mice treated with a vector encoding GFP (n = 10; black circles). Exendin-4 treated mice gained slightly less weight relative to GFP-treated mice control over the 12-week test period [*P* = 0.04]. PYY-treated mice also gained slightly less weight than GFP-treated mice. However, this difference was only apparent during the latter weeks of testing [*P* = 0.0004, *interaction*]. Mice treated with a vector encoding both exendin-4 and PYY gained sustainably less weight relative to GFP-treated mice over the 12-week testing period [*P* = 0.00004]. Data are presented ± SEM.

Of the stimuli tested, the responses of vector treated WT mice to intralipid were the most affected. As such, we then speculated that the responses of these mice were affected by changes in their ability to detect the fatty acids that comprise intralipid. When sampling LA test stimuli after being conditioned to avoid a 1% solution of linoleic acid, WT mice readily learned this aversion as indicated by an observed significant main effect of genotype [*F*(1,6) **=** 10.14, *P* = 0.02;; Fig. 6A], as well as a significant concentration by genotype interaction [*F*(2,12) **=** 6.40, *P* = 0.01]. Posthoc testing revealed that relative to saline treated controls, these WT mice significantly suppressed their lick rate when sampling both 1 and 2% LA [both *ts*(6) ≥ 3.16, *P* ≤ 0.02]. However, after conditioning, relative to saline treated controls, PYY KO mice did not significantly avoid the LA TS [*P* = 0.36;; Fig. 6A]. No concentration by genotype interaction was observed [P = 0.83]. Interestingly, after conditioning, WT mice also generalized the aversion learned to 1% LA to a 5% solution of intralipid [*t*(6) = 2.83, *P* = 0.03; Fig. 6B]. However, PYY KO mice did not avoid this stimulus [*P* = 0.35]. There were no significant differences between the groups in terms of their body weight, mean water intake during the first 2 d of the conditioning phase, mean licks/trial, and mean number of trials taken during Davis rig training.

**Figure 6.**
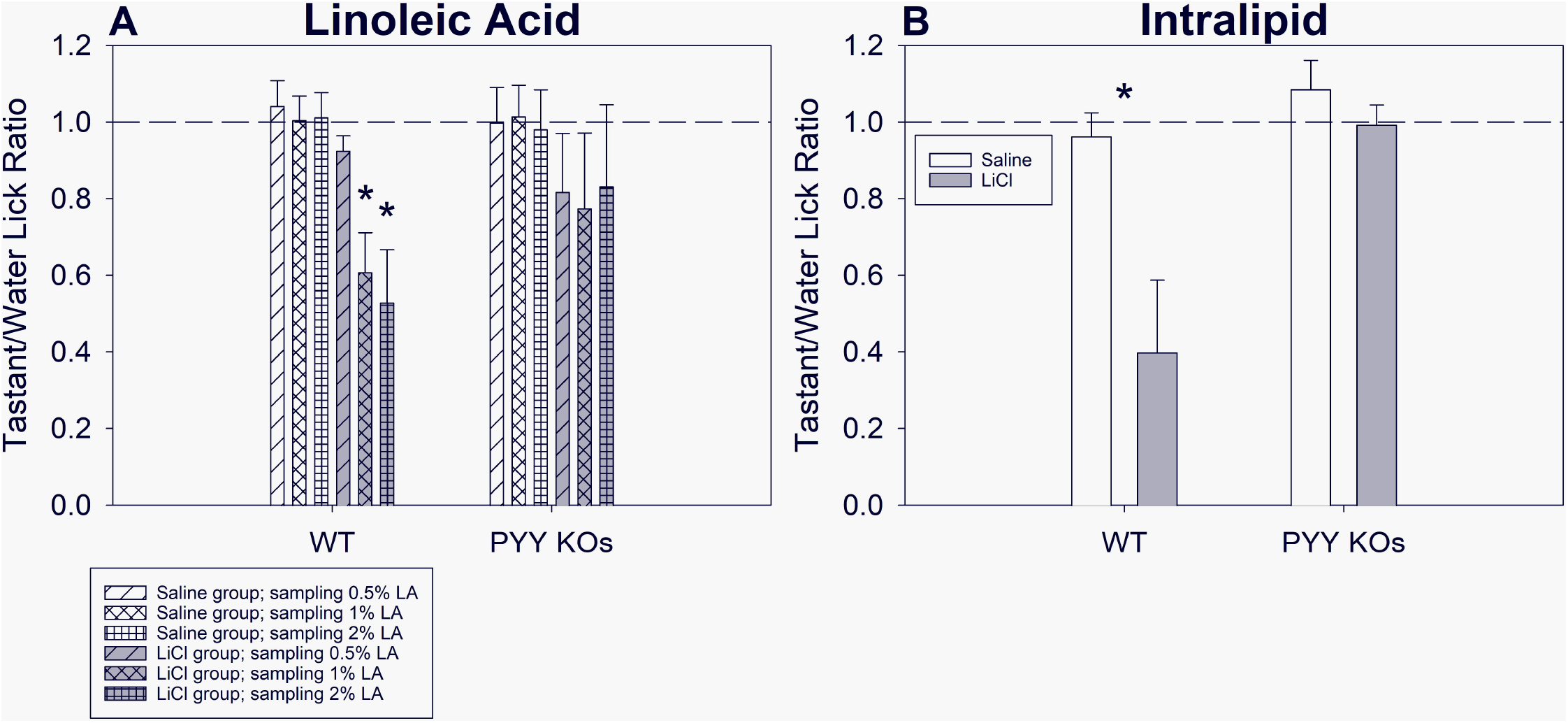
Oral PYY signaling impacts on the ability of mice to learn an aversion to linoleic acid. Brief-access taste testing of WT mice treated with saline (n = 4); WT mice treated with LiCl (n = 4); PYY KO mice treated with saline (n = 4); and PYY KO mice treated with LiCl (n = 4) in response to (A) LA test stimuli (i.e., 0.5, 1, and 2% LA) and (B) 5% intralipid. WT mice that were conditioned to avoid a 1% solution of linoleic acid significantly avoided both 1 and 2% LA [both *Ps* = 0.02] and readily generalized that aversion to a 5% solution of intralipid [*P* = 0.03]. However, PYY KO mice did not significantly avoid any of the TS. The average number of licks per trial for each concentration was divided by that animal’s average licks per trial to vehicle yielding a tastant/vehicle lick ratio. Data are presented ± SEM.

## DISCUSSION

Here we present data suggesting that wild-type mice whose submandibular salivary glands were treated with vectors encoding certain metabolic peptides respond differentially to sapid taste compounds as compared to wild-type mice expressing normal levels of these peptides. When sampling the normally preferred stimulus sucrose, none of the treated mice responded differently than did controls. However, mice treated with vectors encoding both exendin-4 and PYY were hyperresponsive to intralipid relative to GFP treated mice in both the water-deprived and food and water-restricted conditions. Interestingly, when sampling normally avoided stimuli, exendin-4 encoding vector treated mice were hyporesponsive when sampling NaCl relative to control mice, while those treated with a vector encoding PYY were hyporesponsive when sampling quinine. Both of these effects, while significant, were relatively minor in magnitude.

Douton *et al*., investigated the influence of exendin-4 on taste-related behaviors in rats. However, this was done using systemic peptide delivery (i.e., intraperitoneal (IP) injection). They showed that exendin-4, delivered systemically, failed to influence taste-related appetitive reactions towards intraorally infused sucrose in rats (Douton, Norgren, & Grigson, 2021). However, exendin-4 treatment did decrease aversive reactions towards intraorally infused quinine. Systemic treatment also led to reduced approach-related behavior, as well as a reduction in the ingestion of both sweet and bitter solutions suggesting that exendin-4 can influence both appetitive and consummately behaviors, likely via distinct mechanisms (Douton, et al., 2021).

We did not observe any impact of exendin-4 vector treatment on taste related behavior towards quinine. It is possible that the differences in the observed results were the function of the different species used in these experiments. However, we speculate that these distinctions likely also result from differences in the nature of peptide treatment in these experiments (i.e., systemic vs local).

Using IP delivery, Treesukosol *et al*., also investigated the influence of exendin-4, as well as cholecystokinin (CCK), on taste-related behaviors in rats (Treesukosol & Moran, 2022). The authors observed a decrease in the number of trials initiated, as well as a decrease in unconditioned lick responsiveness towards both sucrose and intralipid. Supplementation with CCK did not impact on behavior greater than that observed when exendin-4 was administered alone. Similarly, in a different report, IP treatment with exendin-4 was shown to affect the reinforcing strength and consumption of a highly palatable sweet/fat stimulus in rats (Bernosky-Smith, et al., 2016).

In the current report, we did not observe any impact of exendin-4 vector treatment on taste related behavior towards sucrose or intralipid. Again, we speculate that these differences may be species related or perhaps more likely they result from central effects of systemic peptide delivery. More research is needed to tease apart local vs central impacts of metabolic peptide signaling on the unconditioned licking behavior elicited by appetitive compounds.

We have previously shown that peptide signaling can impact on unconditioned licking behavior towards various gustatory active compounds (i.e., Crosson, et al., 2019; Dotson, Geraedts, & Munger, 2013; Elson, et al., 2010; Hurtado, et al., 2013; La Sala, et al., 2013; Malone, et al., 2021; B. Martin, et al., 2009; Shin, et al., 2008). We hypothesize that these changes in the response to these compounds is rooted in changes in the ability of affected mice, whether wild-type or KOs, to detect these taste compounds. To test this hypothesis, we conducted CTA aversion experiments in wildtype and PYY KO mice. Wild-type mice readily learned to avoid a 1% solution of LA, the major component fatty acid in intralipid, were as PYY KOs were refractory to learning this aversion. As a function of this lack of learning, the KO mice also did not avoid intralipid when sampling after conditioning. Collectively, these data suggest that peptide signaling impacts the ability of mice to detect the fatty acid components making up intralipid, and, as such, impacts on their unconditioned responsiveness to this compound. These mice were global KOs and not vectored treated mice. As such, it is possible that the loss of peptide-mediated signaling in extra-oral tissues of the germline KOs may be responsible for our observed results. Indeed, while our CTA results are suggestive of a detection deficiency in PYY KO mice, alternative explanations still exist (e.g., memory deficits). The use of tissue-specific KOs and/or assessment of these animals with more sophisticated psychophysical measures (e.g., operant conditioning procedures) would be needed to fully disentangle these competing hypotheses.

It is well known that systemic or direct CNS treatment with GLP-1 analogs significantly decrease food intake and body weight in both humans and rodents (e.g., Astrup, et al., 2012; Barreto-Vianna, Aguila, & Mandarim-de-Lacerda, 2016; Hayes, Kanoski, Alhadeff, & Grill, 2011; Mack, et al., 2006; Nayak, Govindan, Baskar, Kalupahana, & Singh, 2010; Szayna, et al., 2000; Yang, et al., 2017; Yang, Moghadam, Cordner, Liang, & Moran, 2014a). Our results suggest that oral expression of exendin-4 is insufficient to impact upon body weight gain. Indeed, a previous report suggests that systemic expression of exendin-4 in salivary glands via AAV5 leads to the detection of active exendin-4 in blood, as well as a significant decrease in weight gain in both mice and rats (Di Pasquale, et al., 2012). The ability to increase circulating levels of peptides via the infection of salivary glands, which can secrete their products via both endocrine and exocrine pathways, is well documented (e.g., Baum, Alevizos, Chiorini, Cotrim, & Zheng, 2015; Borkent & Moharamzadeh, 2017; Di Pasquale, et al., 2020; Lodde, et al., 2006; Voutetakis, et al., 2005; Wang, Wang, Xu, Ding, & Guo, 2014; Zolotukhin, 2012). In previous experiments, to increase the specificity of the AAV5 vector for salivary gland transduction and thereby decrease systemic circulation of peptides from viral infection, we included micro RNA target sites for miR122 and miR206, which are liver and skeletal muscle specific, respectively (Crosson, et al., 2019; Geisler, et al., 2013). Using this micro RNA target site containing vector, we demonstrated that we were able to eliminate leaky expression in the liver and, as a result, were able to keep circulating levels of peptide significantly lower than seen in wildtype mice (Crosson, et al., 2019).

By treating mice with vectors encoding both PYY and exendin-4 and thus causing expression of these products into saliva and not in plasma, we observed a significant attenuation in body weight gain. These results suggest that local peptide signaling in the oral cavity can result in changes in responsiveness to taste compounds, that lead to changes in food intake and subsequent body mass accumulation. However, additional data is needed to confirm this hypothesis.

Investigating the influence of taste modulation on feeding behavior and wholebody metabolic regulation, and meal pattern analysis (a detailed analysis of the size, frequency, and temporal distribution of meals in free feeding animals) would allow for an examination of exactly how ingestion occurs. In addition, metabolic phenotyping in these animals would provide additional information related to energy expenditure, respiratory exchange rate, and locomotor activity. Lastly, while it is posited that the change in body mass accumulation seen in the animals is due to changes in the functioning of peripheral gustatory tissues, it is possible that these changes are mediated by alterations in signaling from other sensory systems present in the oral cavity (e.g., somatosensory, olfactory). Exploring these questions will allow for an ascertainment of the contribution of the gustatory system in the regulation of body mass, and potentially provide an additional physiological target in the development of selective pharmacological interventions targeting pathways designed to help curb hyperphagia and promote body mass regulation and normalization.

## ACKNOWLEDGMENTS

This work was supported by the National Institutes of Health [grant numbers: P30-DC010763, R01-DK62302 to S.Z., and R01-DC012819 to C.D.D and S.D.M.].

All data used is available and the senior author has full access to the data reported in the manuscript.

Authors Zolotukhin and Dotson have filed for a patent entitled “Satiation peptides for weight loss and altered taste sensitivity” (US20170326257A1).

